# Grazing hinders seed dispersal during crop failure in a declining oak woodland

**DOI:** 10.1101/2023.04.14.536763

**Authors:** Pedro G. Vaz, Miguel N. Bugalho, Jose M. Fedriani

## Abstract

Masting, the synchronized production of variable quantities of seeds, occurs in many grazed systems and can be interspersed with years of extreme crop failure, whose frequency and unpredictability are increasing with climate change. Yet, the combined impact of crop failure and grazing on seed dispersal and seed-to-seedling transition remains poorly understood. To address this concern, we investigated rodent-mediated cork-oak (*Quercus suber*) acorn predation, dispersal, and seedling emergence in cattle grazed and non-grazed areas in central Portugal during years with contrasting masting seasons. We found that extreme crop failure led to six times longer and faster acorn dispersal, with 83% more dispersal events than during a year of reproductive success. The percentage of predated acorns also increased by 84%. However, the higher acorn predation was offset by a 2.4-fold higher percentage of unpredated dispersed acorns recruiting into seedlings. Both years ended up recruiting a similar number of seedlings. Acorns emerged seedlings 3.4 times farther in the crop failure year than in the crop success year. Cattle grazing was the main constraint on seed dispersal distance, reducing it by 51% during the extreme crop failure year, while having no noticeable effect during the successful crop year. Our study provides empirical evidence that cattle grazing modulates how an extreme crop failure year can surprisingly be an opportunity for trees remaining fecund to have seedlings established farther apart than in a crop success year. If we are to better manage and preserve the high conservation and socio-economic value of Mediterranean cork oak woodlands in the face of climate change, we must prioritize fecund trees and carefully manage seed dispersal factors such as cattle grazing, particularly during years of crop failure.

## 1. INTRODUCTION

The reproductive strategy of many perennial plants consists of highly variable and synchronized inter-annual seed production (i.e., masting; Kelly, 1994). The benefits of masting have been explained by well-supported hypotheses relating to economies of scale of seed production (Pearse et al., 2020). Predator satiation is one of the most prominent explanations, stating that mast years enhance pre-dispersal seed survival by satiating seed predators (Silvertown, 1980; Kelly and Sork, 2002; Huang et al., 2021; Zwolak et al., 2022). However, there is little empirical data on intermast crop failure years (CFYs), despite its potential consequences for plant regeneration (Koenig, 2021). While climate change is making extreme CFYs more common or stochastic (Hanley et al., 2019; Bogdziewicz et al.; 2020a, 2021; Bogdziewicz, 2022), there is a need to better understand seed predation, dispersal, and seedling establishment in these years compared to crop success years (CSYs). This knowledge is particularly important in Mediterranean climates where masting dynamics is changing under progressive warming and acorn crop failures in oak woodlands are increasing (Fernandez-Martinez et al., 2012; Pérez-Ramos et al., 2015; Chemnitz, 2019).

Cork oak (*Quercus suber*) Mediterranean woodlands are multiple use silvopastoral systems with masting cycles, where animal-mediated acorn dispersal is critical to the ecosystem sustainability (Pons & Pausas, 2006; Aronson et al., 2009; Campos et al., 2013). Cork oak woodlands are classified as “high nature value farming systems” and have high conservation and socio-economic value (Paracchini et al., 2008; Bugalho et al., 2011). In the last decades, a lack of regeneration and increased oak mortality have been reported, which may be endangering the ecological sustainability of cork oak woodlands (Brasier, 1992; Plieninger et al., 2010; Fedriani et al., 2017; Vaz et al., 2019). Like in many plant populations, oak recruitment depends on multiple factors including acorn crop size, predation, dispersal, and seedling recruitment and establishment (Pulido and Díaz, 2005). The oak woodland understory, usually grazed by ungulates, is dominated by grasslands and shrubs. In the Euro-Mediterranean, acorn dispersal is often carried out by rodents and by the European jay (*Garrulus glandarius*), although the latter is less common or absent in cork oak woodlands with low tree density (Herrera 1995; Pons & Pausas, 2008).

Grazing by wild and domestic ungulates is likely to affect seed availability, seed predation and dispersal, and diminish tree-seedling establishment (Levey et al., 2002; Pulido & Díaz, 2005; Leal et al., 2022). Yet, the consequences of ungulate grazing for animal-mediated seed dispersal in contrasting CSYs and CFYs have been overlooked. Research shows that scatter-hoarding rodents and their habitat are affected by the presence of large ungulates (Smit et al., 2001; Torre et al., 2007; Bush et al., 2012; Navarro-Castilla et al., 2017), but how grazing impacts on distance and time to rodent- mediated seed dispersal remains largely unknown (Muñoz & Bonal, 2007). Both short and long seed dispersal distance events are key to the recruitment process and thus for the population persistence and expansion (e.g., Clark et al., 1999; Seget et al., 2022). Also, the time mediated between seed shedding and dispersal is critical to its survival, determining pre-dispersal exposure to predators and pathogens, desiccation, and, ultimately, seed death (Hadj-Chikh et al., 1996; Walter et al., 2013). Particularly, there is a lack of empirical data comparing seed dispersal in grazed and ungrazed areas.

Some studies have emphasized a suite of driving factors on rodent-mediated seed dispersal in oak-dominated systems with masting reproduction, where these animals are key and pervasive seed dispersers. Very few studies, however, compared the driver factors between CSYs and CFYs (Puerta-Piñero et al., 2010). The microhabitat where the acorn is prior to dispersal, in open areas or underneath shrubs, can be of primary importance for the dispersal (Pérez-Ramos & Marañón, 2008; Garcia-Hernández et al., 2016). The microhabitat of acorn arrival (open or shrub) is also a potential explanatory variable of dispersal distance and fate (Perea et al., 2011a; González-Rodríguez & Villar, 2012). Acorns are major food resources for rodents and both distance and time to dispersal are likely to be influenced by the acorn size (Gómez et al., 2008; Morán-López et al., 2018). As rodent abundance influences the number of interactions with acorns, rodent density is also key for acorn predation and dispersal (e.g., Sunyer et al., 2013, 2014; Navarro-Castilla et al., 2017).

In this study, we examine the integration of masting and grazing effects by tracking cork-oak acorns from dispersal to seedling emergence in grazed and non-grazed areas of central Portugal during years of successful and failed crops. Our empirical approach focuses on understanding rodent-mediated seed dispersal during extreme acorn CFYs.

We also investigate the effects of livestock grazing on distance and time to acorn dispersal through comparison with long-term grazing exclosures. We addressed the following questions and hypotheses:

1. How does rodent-mediated acorn dispersal (distance and time to event) differ between CSYs and CFYs? We hypothesized that in a CFY, when acorns are scarce and rodents should quickly move them farther away from conspecifics, the time to dispersal would be shorter and the dispersal distance longer.
2. Does cattle grazing influence rodent-mediated acorn dispersal? We hypothesized that grazing, which can reduce habitat quality and increase rodent stress, would negatively affect acorn dispersal distance.
3. How does rodent density influence acorn dispersal? We hypothesized that higher rodent densities would increase the likelihood of acorn encounters, resulting in shorter distances and times to dispersal.
4. What are the effects of acorn size (weight) and microhabitat type (open areas or underneath shrubs) on rodent-mediated dispersal? We hypothesized that heavier acorns would disperse first and farther. Furthermore, considering rodents’ preference for shrubs in Mediterranean oak woodlands, we expect faster and closer dispersal of acorns underneath shrubs compared to those in open microsites.

## 2. MATERIALS AND METHODS

### 2.1. Field methods

#### 2.1.1. Study area

The study was conducted in central Portugal (38°49’49.526 “N 8°49’15.899 ”W), in a 11000 ha dominated by cork-oak woodlands (Fig. 1) known in Portugal as *montados* (*dehesas* in Spain) within the state-owned farm Companhia das Lezírias (CL). Since the XIX century, CL has been managing cork-oak *montados* for cork harvesting, cattle raising, hunting, and, lately, for biodiversity conservation. Different CL *montado* areas have been fenced to exclude grazing by cattle and to foster oak natural regeneration. Grazing exclusion provides an opportunity to explore the interactions between grazing and rodent-mediated acorn dispersal overtime.

**Figure 1.**
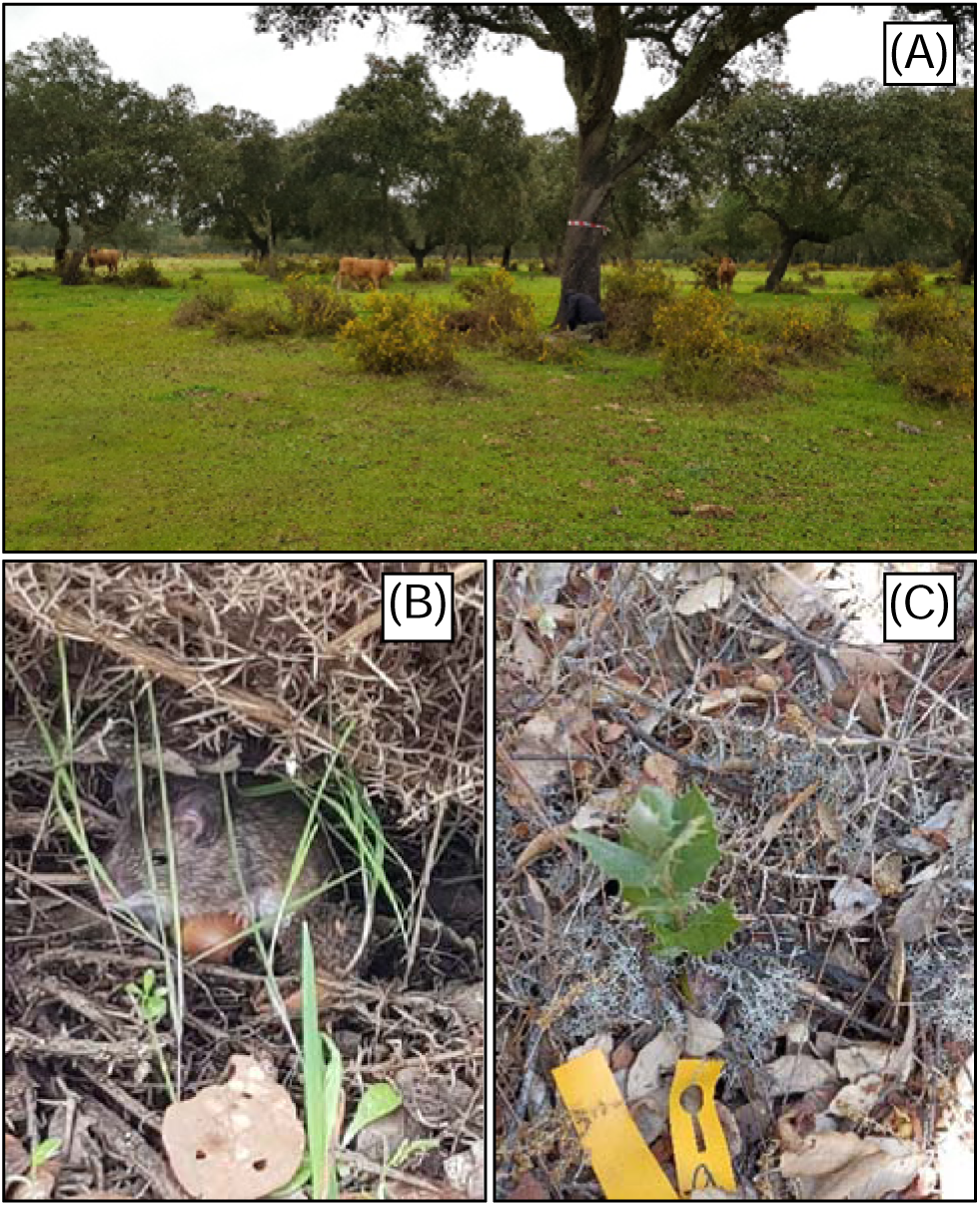
Cork oak (*Quercus suber*) woodland and an acorn-dispersing rodent. (A) grazed study site; (B) scatter-hoarding wood mouse (*Apodemus sylvaticus*); and (C) seedling established from a rodent-dispersed acorn.

The climate in the study area is Mediterranean, with hot, dry summers and cool, wet winters. Mean annual precipitation is 608 mm and mean annual temperature is 16 °C. The slopes are mild, and the altitude range from 1 to 53 m. Soils are well-drained deep Haplic Arenosols (WRB, 2006) with low soil water retention capacity. Cork-oak trees in CL have 24–49 % canopy cover (Listopad et al., 2018). Cork-oak acorns usually fall from late September to early December. Understory vegetation is sparse shrubland dominated by *Ulex australis*, *Cistus salviifolius*, *Myrtus communis*, *C. crispus*, *Genista triacanthus*, and *C. ladanifer*. The grassland understory in open areas include annual species, dominated by the forbs *Tolpis barbata* and *Plantago bellardii*, the graminoids *Agrostis pourretii* and *Avena barbata*, and the legumes *Ornithopus compressus* and *Trifolium arvense*. The wood mouse (*Apodemus sylvaticus*) and the Algerian mouse (*Mus spretus*) are the main acorn dispersal rodents in the sites (see Gonçalves et al., 2013) although they also depredate some acorns. The main acorn predators in CL are year-round grazing wild boars (*Sus scrofa*), Iberian pigs (*Sus scrofa domesticus*) grazing freely outside the grazing exclosures between October and December, and sporadically fallow deer (*Dama dama*).

#### 2.1.2. Experimental design

To evaluate the effects of grazing on acorn dispersal distance and time to dispersal (hereafter acorn-dispersal), we selected nine pairs of sites, each pair including one grazing exclosure site and one cattle-grazed site. The nine ungrazed sites consisted of three sites each that had not been grazed since 1997, 2004, and 2008. Therefore, by the beginning of the experiment, these sites had experienced grazing exclusion for 21, 14, and 10 years, respectively. Across sites, cork-oak trees have a mean basal area of 13.0 m^2^/ha (± 0.6 SE) and a density of 53.6 trees/ha (± 1.2 SE) (Mexia et al., 2022). The distance from any site pair to the nearest one averaged 1191 m (± 488 SE; range = 255– 4720 m). Within site pairs, the mean distance between the grazed and the ungrazed site was 407 m (± 70 SE; range = 218–912 m). Within each site, we haphazardly selected two focal cork-oak trees (≥25 cm DBH) 102 m apart (± 3 SE; range = 93–137 m).

Because long-distance acorn dispersals (e.g., > 30 m) by our target rodent species are likely to be rare (Pons & Pausas, 2007a; Morán-López et al., 2018), we anticipated ∼100 m to ensure independence in the acorn-dispersals between the site’s cork-oak trees. To evaluate the effect of microhabitat on acorn-dispersal, the 36 oaks had at least one shrub of the locally dominant species and an open area with only litter and grasses under the tree canopy. The dominant shrubs used as microhabitat were *Ulex* spp. (mainly *U. australis*) on all but three cork-oaks where the species were *M. communis*, *G. triacanthus*, and *C. ladanifer*. All selected oaks were >50 m from a barbed-wire fence.

To assess the effects of supply (i.e., source) microhabitat (open / shrub) on acorn- dispersal, we placed two pairs of acorn supply stations under the cork-oak canopy, one pair under shrub canopy and the other pair in the open area 1 m away from the shrub. Shrub cover has not changed over the duration of the study around our focal cork-oaks. The paired stations were of two types and were located ∼50 cm apart from each other. To isolate the effect of rodents from other potential consumers, one station was protected from birds and large herbivores with a 32 × 25 × 12 cm wire cage (1.4 cm mesh size) and the other was left unprotected (i.e., incage / outcage). To allow rodents to the interior, the wire cage had two 5 × 5 cm entrances (e.g., Perea et al., 2011a). The cages were firmly attached to the ground, ensuring that they were not displaced during the study.

During November 2018, we collected ∼2000 ripe acorns from CL cork-oaks. In 2019, given the extreme failure of cork-oak acorn production in the study region, we collected ∼2000 ripe acorns from cork-oaks within a 100 km radius of CL on November 5–9. As the natural shedding of acorns coincides with their full maturity, we sought to make the collection match this moment (Merouani et al., 2003). So, we collected acorns on the tree that fell very easily from the cupules or recently fallen on the ground with a homogeneous brown color and no signs of desiccation. To discard weevil-infested acorns, we mixed all the acorns and then floated them in water. For the labeling, we drilled a 1-mm hole in the acorn without damaging the embryo, through which we inserted a 3-cm long wire (0.6 mm diameter). For easy relocation, we attached a 14.5 × 1.6 cm yellow numbered plastic label to the wire. This tagging method does not alter acorn dispersal patterns and yields a high recovery rate (Xiao et al., 2006). The weight of the acorn was then recorded to the nearest 0.01 g, with the labeling corresponding to 6–15% of the total weight.

We placed five labeled acorns on the ground per supply station, totaling 1440 acorns (5 acorns × 2 station types × 2 microhabitats × 36 oaks × 2 years). The tracking of each acorn began the day after the supply. We tracked cork-oak acorns during the 2018–19 and 2019–20 seasons, from November 17 to March 18 in the first year and November 12 to January 31 in the second year. In the first year, we monitored the acorns daily until day 9, then every 4 days until day 25, weekly until day 89, and biweekly until day 119. In the second year, we tracked the acorns on day 1, then every 2 days until day 9, every 4 days until day 25, weekly until day 48, and biweekly until day 78.

#### 2.1.3. Data collection

On every visit, we recorded each acorn dispersal location and any event that occurred on it (e.g., predation, seedling emergence). We searched for acorns within a minimum radius of 50 m around the supply station, adding another 30 m centered on each found acorn. Our detection effort was similar throughout the open areas although we increased search effort for acorns located under shrubs, aiming to compensate for the potentially lower tag detectability in this microhabitat. We measured the distance from the acorn to the supply station (dispersal distance) with a laser meter and a target or with a meter tape, both 1-mm precision. For easy relocation, we marked every acorn location with a labeled stick (30–50 × 0.3 cm) (Xiao et al., 2005). To examine whether the acorn- dispersal would vary with the microhabitat of arrival, we documented if the microsite of acorn deposition on that visit was an open area or under a shrub.

We considered that there was an event if the acorn was moved, gnawed or predated (eaten), cached (e.g., hidden in a log or buried in the ground), or went missing. Intact acorns in the station were considered non-events. Acorns predated while still in the supply stations were omitted from the dispersal distance and time to dispersal analyses. We removed depredated acorns from the study area and assigned them their latest dispersal distance. We deemed an acorn depredated when totally or partially consumed with the embryo damaged (radicle plus plumule). From the first week on, acorns with severely wrinkled skin (dehydrated), very discolored (yellowish), and clearly hollow when carefully pressed were considered non-viable and thus removed.

To evaluate the effect of cork-oak crop size on the acorn-dispersal, we used visual surveys on the 36 cork-oaks prior to significant acorn fall in each late September (Koenig et al., 1994). Visual survey is rapid, effective, and has been validated in predicting acorn production in similar agroforestry systems in the Iberian Peninsula (Carevic et al., 2014). Per cork-oak, we obtained the crop size from the sum of acorn counts for 8 seconds in each orthogonal direction of the crown, totaling 32 seconds. One observer completed all the counting.

To examine the effect of rodent density on acorn-dispersals, we conducted three live- trapping sessions of four consecutive nights per focal cork-oak, before (April 2018), between (May 2019), and after the two-year acorn tracking seasons (January 2020). Per session, we set six Sherman traps (8 × 9 × 23 cm) within a 30 m radius of each focal oak, totaling 216 traps (6 traps × 36 trees). We positioned the six traps ≥10 m apart in variable arrangements, each under a shrub to maximize number of captures and thus to attain most accurate estimates of rodent density (Fedriani, 2005). Trap locations were marked on the shrubs with tape and geo-referenced using a GPS unit with 0.3–1 m precision by post-processing. Each trap was set on the first day, baited with a bread toast (4 x 4 cm) with peanut butter, and checked every day at sunrise.

We marked the captured mice individually with a permanent tattoo on their tails (Chen et al., 2016) and released them at their respective capture locations. For marking, we kept the animal restrained without anesthesia within a thick transparent and well aired bag, with only the tail exposed for tattooing. Each tattoo was done using 1-mL syringes and 30-gauge needles replaced between animals and typically took less than 3 minutes. We used green and red human tattooing inks from Killer Ink Limited (Liverpool, UK). The whole procedure was previously authorized (License 332/2018/CAPT) and reviewed by the animal care committee of the Portuguese Institute for Nature and Forest Conservation (ICNF - Instituto de Conservação da Natureza e das Florestas). Mouse capture and handling were in compliance with Directive 2010/63/EU on the protection of animals used for scientific purposes.

### 2.2. Statistical analyses

#### 2.2.1 Dispersal distance

To evaluate the effects of the explanatory variables on the dispersal distance, we conducted generalized mixed-effects modeling in a Bayesian framework. We created the Bayesian models in Stan computational framework (http://mc-stan.org/) accessed with *brms* package in R (Bürkner, 2017). Per acorn, we analyzed the final distance to the supply station. Our exploratory analyses revealed that (i) the distribution of the dispersal distance as the response variable would be strikingly different per year, including different conditional distributions (see below); (ii) different random effect structures would be required per year; (iii) grazing, a major variable of interest to us, would have no noticeable effect on dispersal distance in one year and would have a strong effect in the other. Also, crop size was an explanatory variable in the CSY (2018–19), but not in the CFY (2019–20), when it was zero in all focal trees. For these reasons, we modeled the two years separately, using two generalized mixed-effects Bayesian models.

To assess the inter-annual differences in the dispersal distance, we used the Mann- Whitney test. Specifically, we tested the null hypothesis that the probability of a 2019– 20 acorn dispersing farther than a 2018–19 acorn would be 50%. To evaluate the difference in the distance distributions between years, we computed the kernel density estimates and the bootstrap version of the Kolmogorov-Smirnov test, which corrects for ties and discontinuities in the distributions (Sekhon, 2011).

The two generalized mixed-effects Bayesian models to evaluate the effects on the dispersal distance were defined as follows. We modeled distances as right-censored in the cases of the acorns that went missing, using the *cens* function in *brms*. Since dispersal could eventually occur after the experiment on acorns remaining at the station for the entire time (cached or as non-events), we also modeled them as having right- censored distances. Because dispersal distance was continuously distributed and greater than zero, we used the log-normal family distribution for the 2018–19 model and the gamma family for the 2019–20 model, both with identity links.

As dispersal distance data were nested by supply station, which were arranged in pairs (incage, outcage), and these in turn were nested by tree, we considered using the three as random factors following that hierarchical order. However, the expected log- predictive density leave-one-out cross-validation difference (Δ ELPD LOO) suggested adding station pair to random effects did not improve the expected predictive accuracy of the 2018–19 model (Δ ELPD LOO = 0.0 ± 1.7 SE). Thus, the random part of the 2018–19 model included station, while that of the 2019–20 model included all three factors (tree/station pair/station identifiers). The numeric fixed covariates were acorn weight, rodent density, plus crop size in 2018–19.

In both models, the fixed effects included the categorical variables grazing (levels = grazed, 21-, 14-, 10-year exclosure), station type (incage, outcage), supply microhabitat (open, shrub), and arrival microhabitat (open, shrub). Because preliminary exploratory analysis showed the univariate model with grazing having four levels worsened (Δ ELPD LOO = 2.0 ± 1.6 SE) or equaled (Δ ELPD LOO = 0.8 ± 1.8 SE) the predictive accuracy of the 2018–19 and 2019–20 null models, we opted for grazing with two levels (ungrazed, grazed). The correlation matrices by year for the initial explanatory variables revealed no collinearity problems (≤ 0.63 in all cases).

The minimal adequate (optimal) models were arrived at by first fitting the full models (using all explanatory variables simultaneously) followed by backward elimination of one explanatory variable at a time. We used Watanabe-Akaike Information Criterion (WAIC; Watanabe, 2010) to compare the relative fit of the models to the data, interpreting WAIC differences greater than twice its corresponding standard error as suggesting that the model with the lower WAIC fitted the data substantially better (Vaz et al., 2021). In 2018–19, rodent density and grazing were dropped in that order using the WAIC criterion to reach the optimal model with station type, arrival microhabitat, crop size, and acorn weight. In 2019–20, the dropping order was rodent density and station type, to reach the model with grazing, supply microhabitat, arrival microhabitat, and acorn weight.

To improve convergence while controlling against overfitting, we assigned weakly informative priors to all the effect size beta parameters of the two models (see Gelman, 2022). For the 2018–19 model, we used the *normal (0,3)* distribution for the beta in all levels of categorical variables and the *normal (0,2)* for the beta in the numeric variables except acorn weight wherein we used the *normal (0,1)*. For 2019–20, we used the *normal (0,3)* distribution for the beta parameters in all the variables except for rodent density and acorn weight, wherein we used *normal (0,2)* and *normal (0,1)*, respectively. For each model, we ran four parallel MCMC chains until convergence was reached (all Rhat ≤ 1.1). Each chain had 4000 iterations (warmup = 1000, thin = 1), totaling 12000 post-warmup samples. We assessed model adequacy using posterior predictive checks. Prior to all the analyses, we centered and standardized the numeric covariates. We performed all analyses in R v. 4.1.1 (R Core Team, 2021).

Through Bayesian statistical inference, we estimated the β parameters of the models, their credible intervals, and tested statistical hypotheses (e.g., McElreath, 2018). As the Bayesian inference returns a distribution of possible effect values (the posterior), the credible interval is the range containing a particular percentage of probable values (e.g., 95%). After we have computed the posterior distribution for β, we have interrogated it directly. For instance, if 99% of the posterior distribution is above 0, we were 99% confident that β >0, P(β >0) = 0.99, and 1% confident that β<0 (Azedo et al., 2022).

#### 2.2.2 Time to dispersal

To investigate the association between time to acorn-dispersal and the set of explanatory variables, we used Cox proportional hazards frailty modeling (Therneau and Grambsch, 2000). The response variable was the number of follow-up days until the dispersal event and was modeled as right-censored due to the uncertainty that the dispersal could eventually occur in some acorns after the experiment ended. Because the distributions of the response variable were remarkably different by year and the 2018–19 model had crop size as an additional predictor, we ran a model by year. We used the same full set of fixed covariates by year as in the two full Bayesian models for dispersal distance. However, the correlation matrices by year revealed supply microhabitat was strongly correlated with arrival microhabitat in 2018 (*r*_φ_ = 0.82) and was excluded from this year’s Cox models to prevent collinearity. The effect of supply station was accounted for by including it into both models as a random proportionality factor (“frailty”). Adding station pair to the random effects would have significantly worsened both the 2018–19 (*χ*^2^ = 417.05, P < 0.0001) and 2019–20 (*χ*^2^ = 261.18, P < 0.0001) Cox model fits (*anova* command in the *coxme* package in R; Therneau, 2020).

To fit the two Cox models, we used the *survival* package in R (Therneau, 2021). The significance (chi-square test; P < 0.05) of each predictor was evaluated by backward-stepwise elimination from the full model (Therneau & Grambsch, 2000). In 2018–19, crop size, acorn weight, grazing, and station type were sequentially dropped to reach the optimal model with arrival microhabitat and rodent density. In 2019–20, the dropping sequence was supply microhabitat, acorn weight, station type, and rodent density, reaching the optimal model with grazing and arrival microhabitat. To infer the expected time of acorn first dispersal with estimates of uncertainty by covariate from the two Cox models, we used the *coxed* package in R, via the nonparametric step-function approach (Kropko & Harden, 2020).

#### 2.2.3 Rodent density

To calculate the rodent density (ha^-1^) per year associated to each of the 36 cork-oaks, we used the multi-session dataset from the livetrappings. We used spatially explicit capture-recapture models (Efford, 2004; Borchers & Efford, 2008) fitted via the maximum likelihood method using the *secr* package in R (Efford, 2022a). We used the half-normal detection function for the detection probability. Half-normal has *g0* (encounter rate) and σ (scale for *g0* decreasing with distance to the home-range center) parameters, defining the detection as a function of location. To model *g0*, we ran six models considering the individual effects (learned or transient responses) of animals, traps, and interactions of both (for details, see Efford, 2022b). We modeled σ as a constant in all cases. To select among the six models plus the null model (having constant *g0* and σ), we used the Akaike Information Criterion (*AIC* command in *secr*). Following the AIC, we estimated rodent density considering trap-specific responses on the encounter rate (*g0* ∼ *k*; see Efford, 2022b).

## 3. RESULTS

Sixty-seven percent of the 1440 acorns supplied in the two years were dispersed. The mean dispersal distance from the supply station was 5.91 m (median = 2.01 m, range = 0.01–45.78 m). Between the two years with contrasting crop successes, the dispersal distance and its distribution (Fig. 2A) differed significantly (*U* = 38932, p < 0.0001; *D* = 0.57, p < 0.0001). In the CFY (2019–20), the mean dispersal distance (mean = 8.34, median = 4.63, range = 0.10–45.78 m) increased 6 times compared to CSY (2018–19) (1.46, 0.45, 0.02–22.90 m). In 2018–19, 52.4% of the 720 acorns did not disperse while in the CFY only 12.6% of the 720 acorns were not moved. Further, 75% of the acorns dispersed ≤1.05 m only in 2018–19, while that 75^th^ percentile was 12 times higher in 2019–20 (12.53 m). Ten percent of the 2019–20 dispersals were longer than the maximum dispersal in 2018–19.

**Figure 2.**
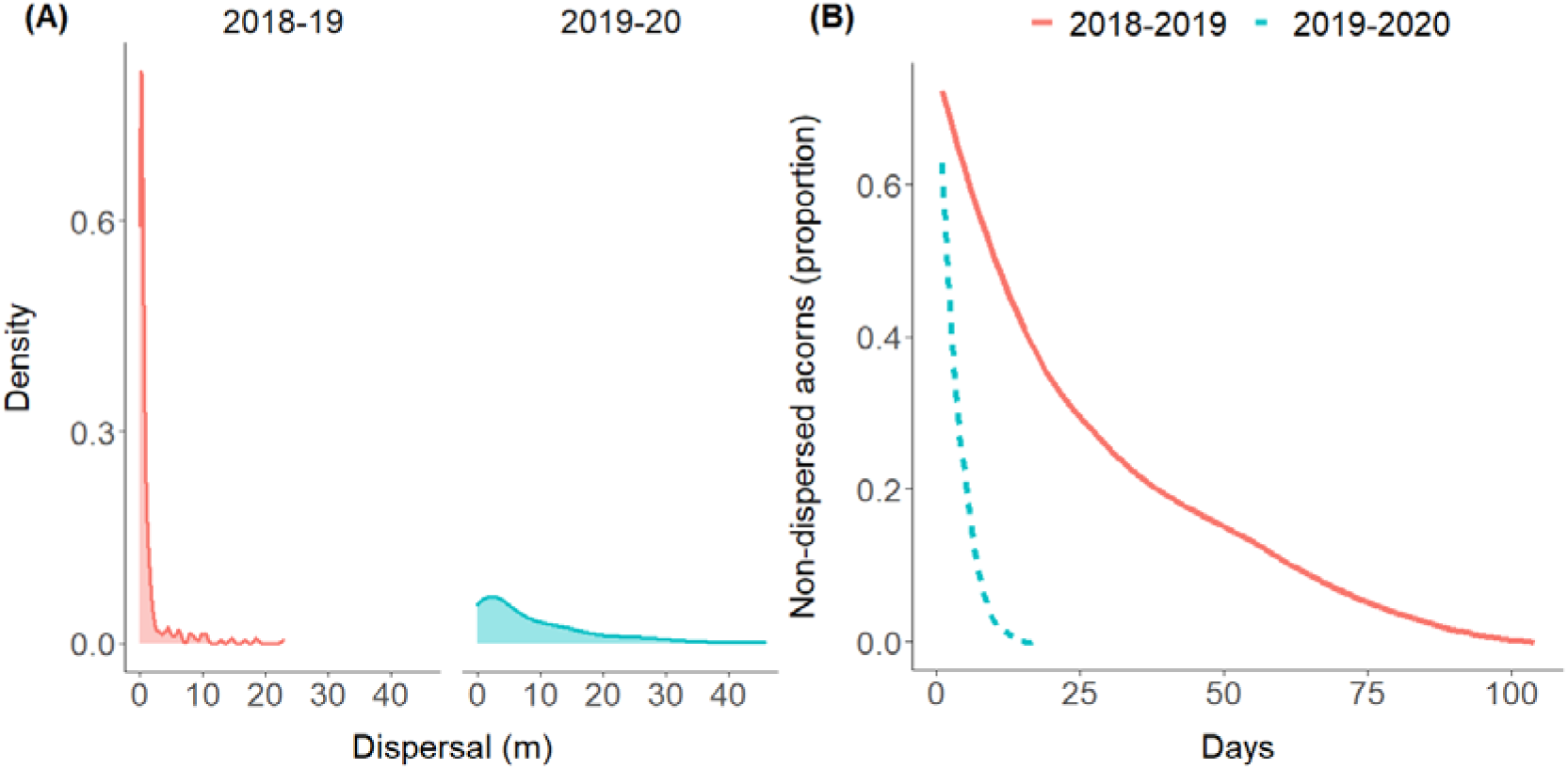
Cork-oak acorn dispersal in years when acorn production was successful (2018–19) and failed (2019–2020). Kernel density of dispersal distance (A); proportion of non-dispersed acorns over time (B).

The first acorn-dispersal occurred on average at day 10 (median 3 days). Between years, the mean duration until dispersal (Fig. 2B) was ∼6 times shorter in the acorn CFY (mean = 4, median = 3 days) than in the CSY (mean = 22, median = 9 days). Also, all acorns dispersed by day 17 in 2019–20, but not until day 104 in 2018–19.

Predation was the most frequent final acorn fate in both years, especially among dispersed acorns (Table 1). Noticeably, 80.8% of the supplied acorns were predated in the second year (cork oak CFY). Seventeen and 13% of acorns went missing in 2018– 19 and 2019–20, respectively. Missing acorns were more frequent for undispersed than for dispersed ones. In both years, barely ∼1% of the overall acorns emerged seedlings. In 2018–19, seven seedlings emerged between February 12 and March 16 (i.e., mean = 49 days after supply; range = 29–104). In 2019–20, eight seedlings emerged between January 14 and 31 (72; 63–78). The percentage of dispersed acorns recruited into seedlings was 7% and 17% in the CSY and CFY, respectively (Table 1). The mean dispersal distance of emerged seedlings was 3.17 (range = 0.27–18.75 m) and 10.76 m (0.40–32.00 m) in the CSY and CFY.

**Table 1.** Final fate of the tracked acorns in years when acorn production was successful (2018–19) and failed (2019–2020).

| Final status |  | Year |  |
| --- | --- | --- | --- |
|  |  | 2018-19<br>% (count) | 2019-20<br>% (count) |
| Dispersed | predated | 33.9 (244) | 80.8 (582) |
|  | cached | 1.5 (11) | 1.5 (11) |
|  | moved only | 7.9 (57) | 0.4 (3) |
|  | missing | 3.3 (24) | 3.5 (25) |
|  | seedling | 1.0 (7) | 1.1 (8) |
|  | *Total* | *47.6 (343)* | *87.4 (629)* |
| Undispersed | predated | 11.5 (83) | 2.8 (20) |
|  | cached | 0.4 (3) | 0.1 (1) |
|  | no event | 26.7 (192) | 0.0 (0) |
|  | missing | 13.8 (99) | 9.7 (70) |
|  | seedling | 0.0 (0) | 0.0 (0) |
|  | *Total* | *52.4 (377)* | *12.6 (91)* |
| *Total* |  | *720* | *720* |

The mean density of dispersing rodents estimated via spatially explicit capture- recapture models on the 36 focal trees was 18 (95% CI =15–22) and 9 (8–11) individuals ha^-1^ in the CSY and CFY, respectively.

### 3.1 Dispersal distance

When we accounted for distance censoring and the effect of random factors, the predictors of dispersal distance differed in the two optimal Bayesian mixed models (Table 2, Fig. 3). Only arrival microhabitat and acorn weight had evident effects on dispersal distance in both years. Mean dispersal distances were 65% (2018–19) and 72% (2019–20) longer in acorns dispersed into open areas than into shrubs. As expected, heavier acorns dispersed farther. For example, a weight increase from 4 to 8 g corresponded to 38% and 46% increases in dispersal distance in 2018–19 and 2019–20, respectively.

**Figure 3.**
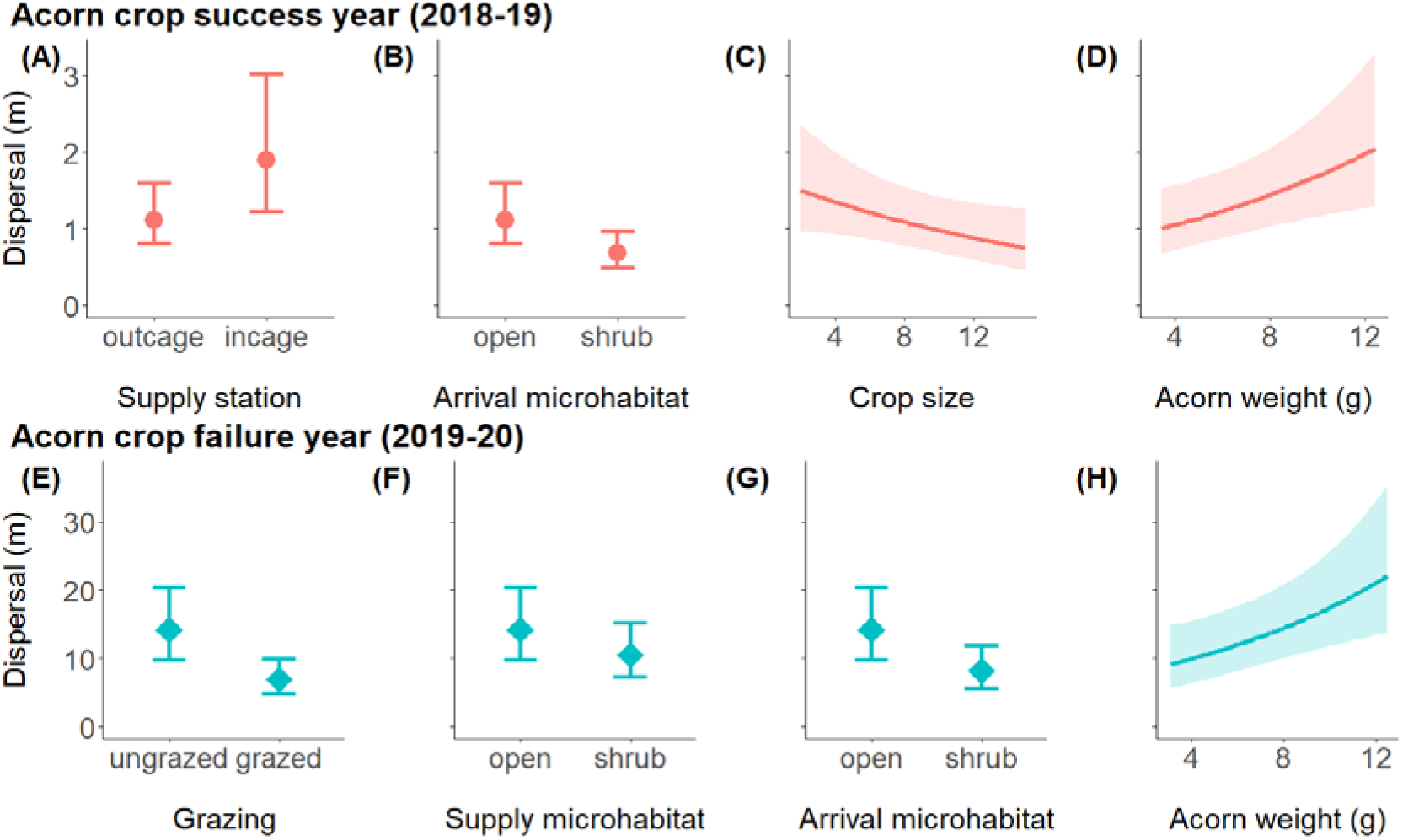
Mean fitted acorn dispersal distances (±95% credible intervals) by grazing and covariates, as predicted by two optimal mixed-effects Bayesian models for when cork-oak acorn production was successful (2018–19; A–D) and failed (2019–2020; E– H). Supply station = type of acorn offering station (inside or outside a wire mesh cage); arrival microhabitat = microhabitat of the latest acorn location (underneath a shrub or in the open area); crop size = acorn count per focal cork-oak (zero on all trees in 2019-20); grazing = cattle grazing; supply microhabitat = microhabitat in which the acorn was offered.

**Table 2.**
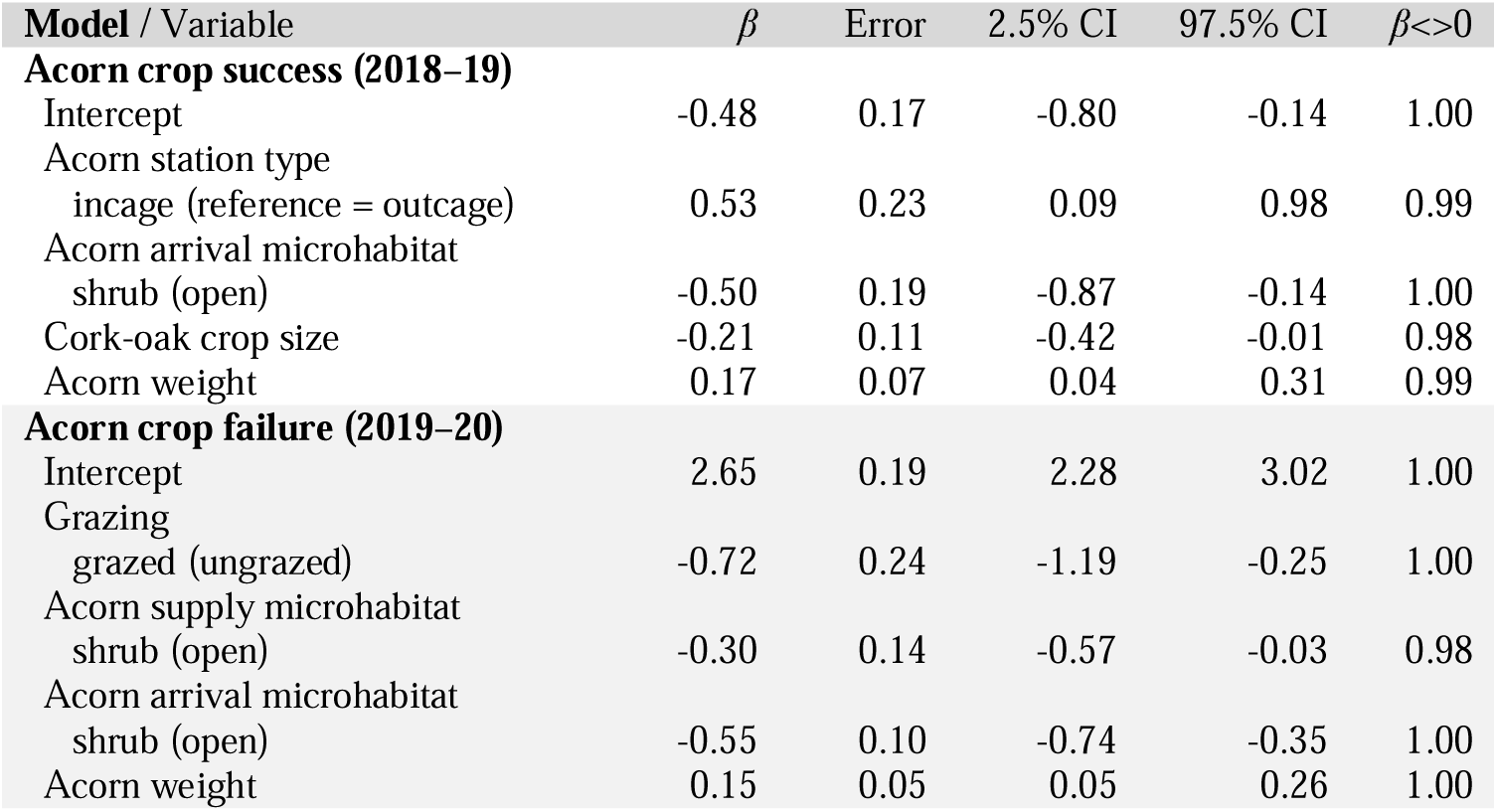
Fixed part of the two optimal mixed-effects Bayesian models predicting the effects of cattle grazing and covariates on the dispersal distance of cork-oak acorns in years when acorn production was successful (2018–19) and failed (2019–2020). CI = credible interval for the β parameter (the range containing 95% of probable values); β<>0 = probability of confidence that the effect is positive (if β > 0) or negative (β < 0). Potential scale reduction factor on split chains (Rhat) was 1.00 in all parameters.

In 2018–19, acorns offered to rodents only (incage supply stations) dispersed 1.7 times farther (posterior distribution mean = 1.89; 95% CI = 1.21–3.02 m) than outcage acorns (1.11; 0.80–1.59). This year, it was also evident that dispersal went farther in trees with smaller crop sizes. For example, tripling the acorn count from 4 to 12 led to 35% shorter dispersal distances. This result was in line with the farther dispersals in the cork oak CFY than in the CSY.

The expected impact of grazing was only evident in 2019–20, the CFY, when it was the variable with the largest negative effect on dispersal distance (Fig. A.1, Appendix 1). The mean of the posterior distribution in 2019–20 was 6.88 m in the grazed sites (95% credible interval = 4.80–9.88), whereas it was 14.12 m in the grazing exclosures (9.80–20.40). Thus, cattle grazing reduced dispersal distance by 51%. This year, acorns supplied in open areas also dispersed 1.3 times farther (14.12; 9.80–20.40 m) than those provided under shrubs (10.49; 7.18–15.11 m).

### 3.2 Time to dispersal

Time to first dispersal had also arrival microhabitat as an important predictor in both years, although with surprisingly opposite effects (Table 3, Fig. 4). Relative to open microsites, shrubs were associated with 71% shorter and 21% longer times in 2018–19 and 2019–20, respectively. The expected negative effect of rodent density on time to first dispersal was evident in 2018–19, particularly up to a density of ∼10 ha^-1^. Last, as with dispersal distance, the effect of grazing on the velocity of first dispersal was only evident in the CFY (2019–20). The mean time to first dispersal was 1.3 times faster at grazed sites (mean = ∼3; 95% CI = 3–4 days) than ungrazed sites (∼5; 4–5 days).

**Figure 4.**
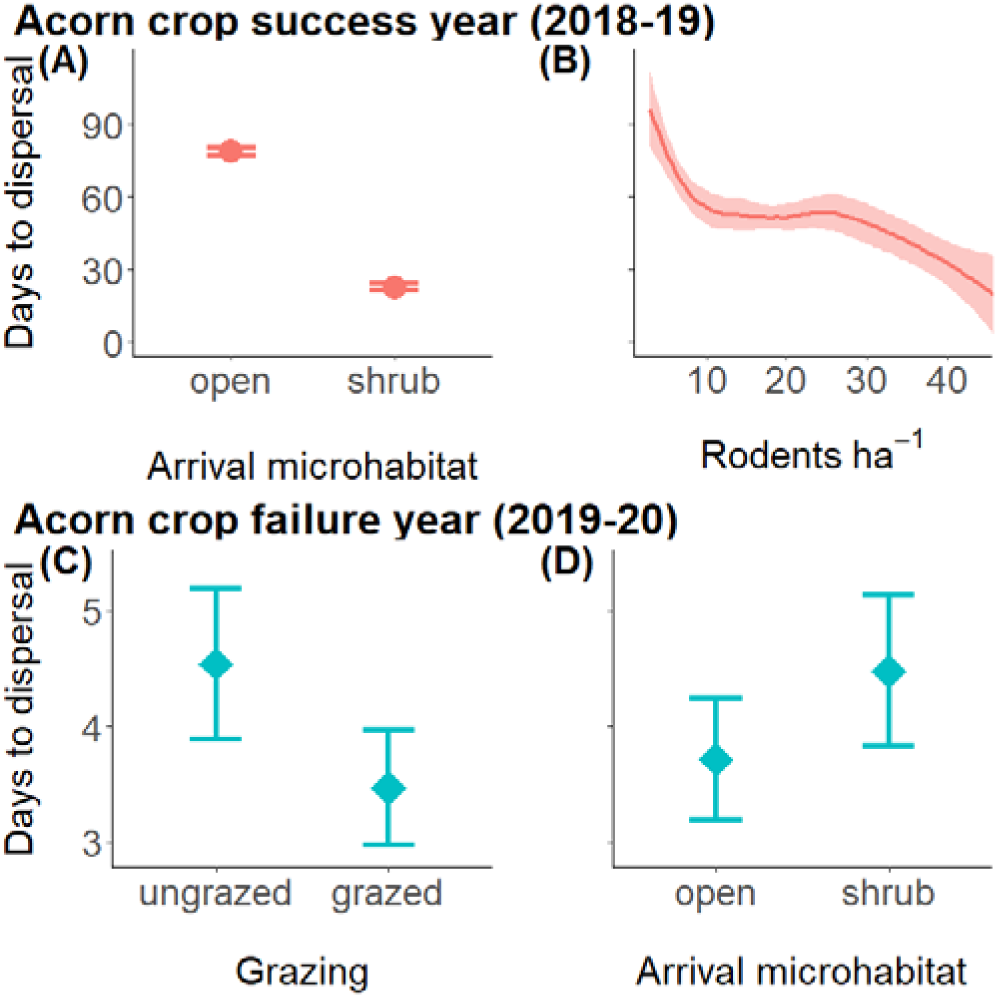
Mean fitted number of days to first acorn-dispersal (±95% confidence intervals calculated via bootstrapping) by explanatory covariates, as predicted by the optimal Cox proportional hazard models with expected durations, when cork-oak acorn production was successful (2018–19; A–B) and failed (2019–2020; C–D). See Fig. 3 for variable descriptions.

**Table 3.** Optimal survival models by Cox multiple regression for relative “risk” of dispersal on cork-oak acorns in years when production was successful (2018–19) and failed (2019–2020). Supply station was used as a random proportionality factor (frailty). HR = hazard rate.

| Model / Variable | $\beta$ | SE | $\chi^2$ | P | HR (95% CI) |
| --- | --- | --- | --- | --- | --- |
| **Acorn crop success (2018–19)** |  |  |  |  |  |
| Acorn arrival microhabitat |  |  |  |  |  |
| shrub (open) | 0.839 | 0.221 | 14.40 | <0.0001 | 2.31 (1.50–3.57) |
| Rodent density | 0.443 | 0.201 | 4.88 | 0.027 | 1.56 (1.05–2.31) |
| frailty |  |  | 975.31 | <0.0001 |  |
| **Acorn crop success (2019–20)** |  |  |  |  |  |
| Grazing |  |  |  |  |  |
| grazed (ungrazed) | 0.460 | 0.163 | 7.99 | 0.005 | 1.58 (1.15–2.18) |
| Acorn arrival microhabitat |  |  |  |  |  |
| shrub (open) | -0.317 | 0.104 | 9.33 | 0.002 | 0.73 (0.59–0.89) |
| frailty |  |  | 473.20 | <0.0001 |  |

## 4. DISCUSSION

Our results provide novel insights on masting dynamics within climate change context and about increasing intermast events of widespread crop failure. Our data show that an extreme CFY can strikingly affect animal-mediated seed dispersal, both through seed scarcity effects and because primary significant covariates may be different or have very different effects from those occurring during a CSY. By combining crop success and grazing effects in a Mediterranean silvopastoral system, we found evidence of cattle grazing effects on the distance and time to cork oak acorn dispersal but only during the extreme CFY. This year had six times longer and faster seed dispersals and 83% more dispersal events, although, consistent with the predator satiation hypothesis, the percentage of predated acorns also increased by 84% as compared to the CSY. Our results suggest predation/dispersal (distance and velocity) trade-offs, as the net outcome for the early cork oak demography was surprisingly similar in such contrasting masting seasons, with only 1% of acorns recruiting into seedlings.

### 4.1 Crop size and failure

Our results showed that acorns in the less productive cork oak trees during the CSY were also dispersed farther. This is consistent with the much longer dispersal distances observed during the CFY (Fig. A.2, Appendix 1). Although past studies on seed dispersal have rarely focused on extreme CFYs, our between-year comparison is in line with research obtained for holm oak (*Quercus ilex*) documenting a less clumped pattern of dispersed seeds in a small crop size year (Puerta-Piñero et al. 2010). Our inter-tree seed dispersal comparison is compatible with few studies documenting shorter dispersal differences with increased seed abundance (Jansen et al., 2004; Moore et al., 2007). As the CSY had simultaneously twice the rodent density and far fewer dispersal events, our results also countered the predator dispersal hypothesis (Vander Wall & Balda, 1977; Vander Wall, 2010), predicting that larger crops may imply higher per capita seed dispersal rates or longer dispersal distances than in non-masting years (Seget et al., 2022). Our data better support the predator satiation hypothesis as a mechanism of mast seeding in cork oak woodlands. This adaptation to predator satiation is common in forest stands dominated by one species (Kelly & Sork, 2002). In mixed species stands with cork oak trees (e.g., mixed cork- and holm-oak woodlands), it may be possible that the availability of other tree species’ seeds may contribute to satiate predators in cork oak CFYs. Some studies have compared seed predation between monospecific and mixed forest stands in masting years, but not in extreme CFYs (Smith, 1987; Hart, 1995; Curran & Webb, 2000).

We suggest that the few acorn-producing cork-oak trees in the extreme CFY may have had more effective dispersals among their acorns escaping predation than in the CSY. This is a counter-intuitive explanation contributing to unveil seed dispersal mechanisms by oak trees in CFYs. We simulated the presence of such fecund trees by supplying acorns in our focal trees in the CFY, although true highly fecund individuals tend to be large, old trees producing more and larger acorns as compared to our focal trees (see Bogdziewicz et al., 2020b). Noticeably, the higher predation in the CFY was offset by a 2.4 times higher percentage (17 vs. 7%) of dispersed acorns recruited into seedlings, with both years ending up recruiting a similar number of seedlings from the supplied acorns. Importantly, seedling emergences occurred ∼3.4 times farther in the CFY than in the CSY. As only 1.5% of the dispersed acorns were cached in each year, caching likely had a negligible effect on decreasing the proportion of predated acorns (Vander Wall, 2002; Xiao et al., 2013, Zwolak et al., 2016).

### 4.2 Grazing and crop failure

Our data showed mixed effects of cattle grazing on acorn dispersal in the extreme CFY, whereas it had no noticeable effect during the CSY. Grazing was the main negative factor affecting seed dispersal distance by reducing it by 51%. Grazing, therefore, seems to attenuate the potential of fecund trees to establish seedlings farther away in CFYs. Promoting regeneration success and oak woodland restoration necessitates the consideration of cattle grazing management during CFYs, namely though the creation of grazing exclusion areas. On the other hand, by reducing the time to dispersal by 30%, grazing had the potential to increase the survival of acorns quickly moved to adequate microsites, since cork oak is a fast-germinating species when conditions are favorable (Amimi et al., 2020). Yet, as shown by a recent meta-analysis of Mediterranean-wide datasets, acorns are much less likely to survive in the presence of ungulates (Leal et al., 2022). It remains possible that an overall low probability of acorn survival in grazed areas, possibly associated with ungulate predation and lower availability of potential recruitment microsites (Clark et al., 1998; Levey et al., 2002; Gómez et al., 2003) in the study area, may obviate the arguable advantage of the higher dispersal speed shown in our data in a CFY. The faster speed of acorn dispersal may further be outweighed by negative impacts of grazing at later oak life stages (Pulido & Díaz, 2005), namely seedling trampling and seedling and sapling browsing (Cierjacks & Hensen, 2004; Muñoz et al., 2009; Parsons et al., 2021). Proportionally to grazing pressure, cattle might also have the potential to indirectly affect acorn dispersal through negative effects on shrub cover (Smit & Verwijmeren, 2011).

### 4.3 Further effects

We also show how the effects of other factors known to make animal-mediated seed dispersal a non-random spatiotemporal process (Pérez-Ramos & Marañón, 2008) can vary from a CSY to a CFY. For example, our hypothesis that acorns underneath shrubs would disperse faster was supported by the result of the arrival microhabitat variable in the CSY, but not in the CFY when acorns were moved 21% faster into open areas than into shrubs. As acorns are high nutritional value resources (Pons & Pausas, 2007a), which are scarce in CFYs, a possible explanation for that result is an attempt by rodents to quickly escape the increased acorn predation pressure by other rodents beneath shrubs (García-Hernández et al., 2016) at the cost of increased risk of their own predation in open grasslands (Sunyer et al., 2016). Similar trade-offs were previously described between higher risks of rodent seed predation beneath vegetation canopies and reduced cache pilferage in open vegetation (Preston & Jacobs, 2005; Muñoz & Bonal, 2011; Steele et al., 2014). On the other hand, given the higher density and foraging activity of small rodents beneath shrubs (Herrera, 1995; Gray et al., 1998; Rosalino et al., 2011) and that most of the acorns in our work were predated, it is not surprising that shorter dispersal distances were found in this microhabitat.

Irrespective of the crop success, larger acorns were dispersed farther away as hypothesized. Some related studies have also established a positive relationship between larger seeds, which offer higher nutritional rewards for rodents than smaller seeds (Smith & Reichman, 1984; Jansen et al., 2002; Vander Wall, 2003), and dispersal distance (Gómez et al., 2008; Puerta-Piñero et al., 2010; Perea et al., 2011b), as well as better seedling performance later on (Seiwa, 2000; Vander Wall, 2001; Gómez, 2004; Moles & Westby, 2004; Cicek & Tilki, 2007). The high individual variation in seed size of many plant species (Herrera, 2009; Wang & Ives 2017) having similar dispersal processes can thus lead to a wide range of rodent-mediated dispersal distances.

Interestingly, as the cork oak is the only oak species with annual and biennial acorns of widely varying sizes on the same tree (Elena-Roselló et al., 1993; Ramos et al., 2013), our results may imply an even more pronounced adaptation of the species to short and long-distance dispersal, both important for the population persistence and expansion. On the other hand, we found no evidence that larger acorns were dispersed more rapidly in the two contrasting crop years, which may indicate that the positive selection, reflected in greater dispersal distances, occurs after the acorns are removed, and less by visual assessment as in the case of the European jay (Pons & Pausas, 2007b).

## 5. Conclusions

Our study provides empirical evidence that extreme CFYs can be opportunities for fecund cork-oak trees to have seedlings established at greater distances and in similar numbers as in a CSY. This phenomenon can be enhanced by greater emergence rates among the fewer acorns escaping predation, associated with more and faster dispersal events. We also revealed that cattle grazing can markedly alter seed dispersal patterns during an extreme CFY, strongly decreasing dispersal distances and increasing the speed of cork oak acorn dispersal, even though it may show no noticeable effect during a CSY. Considering the rising occurrence of acorn crop failures in many oak woodlands and the ongoing decline of Mediterranean oak silvopastoral systems (e.g., cork and holm oak woodlands) partially attributed to overgrazing, the previously unrecognized interplay between grazing and crop failure warrants attention. To safeguard the ecological and socio-economic value of these systems amidst climate change, effective management strategies for seed dispersal drivers, such as grazing during CFYs, should be acknowledged and prioritized. Successfully preserving and restoring high-value oak woodlands requires management decisions that carefully consider the trade-offs between acorn predation and seedling establishment, as well as the impact of shrub cover on acorn dispersal. Despite being a significant challenge, recognizing the influence of these multiple drivers in management decisions is crucial.

## Acknowledgements

We are deeply grateful to the employees of Companhia das Lezírias for the logistical support, especially to Rui Alves and Sandra Alcobia. PGV funded by: Portuguese Science and Technology Foundation (SFRH/BPD/105632/2015; CEABN-InBIO indirect costs [overheads] UID/BIA/50027/2020; cE3c funding UIDB/00329/2020- 2023). We thank Abdullah Ibne, University of Lisbon, and to undergraduate students at Lusófona University, Lisbon, for fieldwork assistance.

