## Appendix A for "Grazing hinders seed dispersal during crop failure in a declining oak woodland": SupportingInformation.pdf

Vaz PG, Bugalho MN, Fedriani JM. 2023. Cattle grazing alters how cork oak crop failure leads to faster and farther rodent-mediated acorn dispersal.

### Appendix A

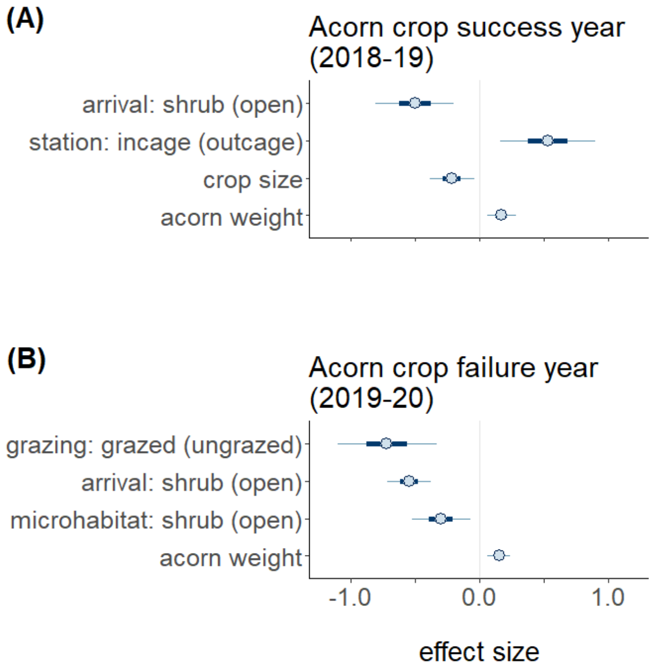


**Figure A.1.** Mean effect sizes (±95% and 50% credible intervals) of cattle grazing and covariates on the dispersal distance of cork oak acorns as predicted by the two optimal mixed-effects Bayesian models fitted in years when acorn production was successful (2018–19) and failed (2019–2020). Arrival = microhabitat of the latest acorn location; station = type of acorn supply station (inside or outside a wire mesh cage); crop size = acorn count per focal cork oak (zero on all trees in 2019-20); grazing = cattle grazing; microhabitat = microhabitat in which the acorn was supplied. The reference level is between parentheses for categorical variables.

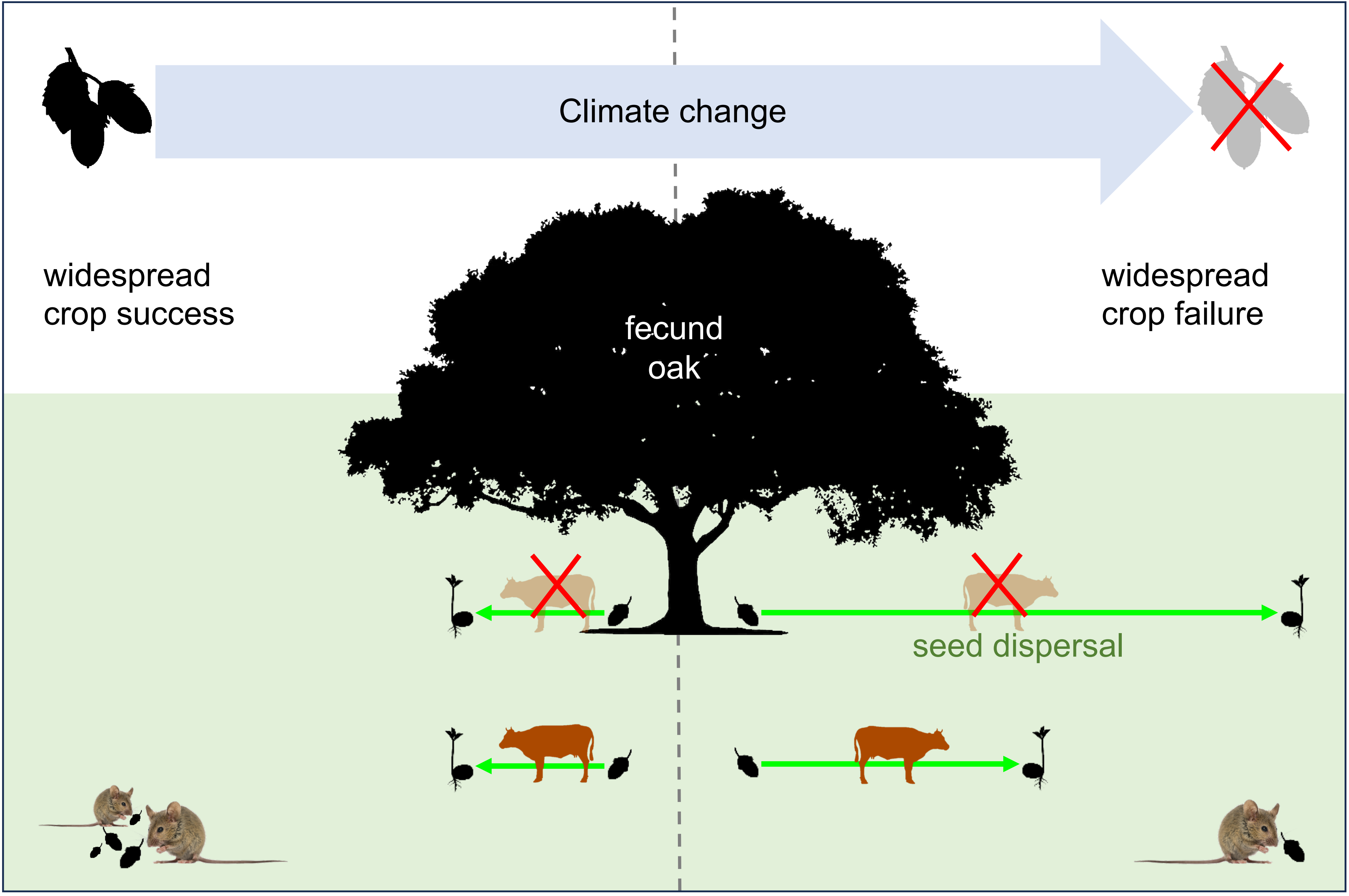

Climate change
widespread
crop success
widespread
crop failure
fecund
oak
seed dispersal
